# Unveiling stem cell induction mechanisms from spatiotemporal cell-type-specific gene regulatory networks in postembryonic root organogensis

**DOI:** 10.1101/2024.06.13.598926

**Authors:** Javier Cabrera, Álvaro Sanchez-Corrionero, Angels de Luis Balaguer, Laura Serrano-Ron, Cristina del Barrio, Pilar Cubas, Pablo Perez-Garcia, Rosangela Sozzani, Miguel Moreno-Risueno

## Abstract

Plants grow continuously by developing new organs, a complex process that requires the formation of specific and functional tissue patterns. Tap root systems, as observed in *Arabidopsis thaliana*, undergo lateral root formation, a developmental mechanism that necessitates the establishment of stem cell lineages. However, the underlying mechanisms remain poorly understood. We have reconstructed a spatiotemporal cell-type-specific transcriptional map of early lateral root organogenesis in Arabidopsis, profiling single and double fluorescent markers across 8 different cell types in the root stem cell lineage. Employing dynamic Bayesian network inference, based on time-course experiments and developmental time, alongside tree-based methods, we investigated lineage developmental progression and precursor stem-cell specification. Our results reveal a morphogenic cascade of hierarchical interdependent transcription factors driving stem cell initiation, and identify the QC/Endodermis transitioning cells as root stem cell progenitors. The associated formative program involves a profound transcriptomic re-arrangement, which, remarkably, precedes the activation of known stem-cell transcriptional signatures. Our data support a model in which root-stem-cell networks do not initiate stem formation, although various stem cell regulators are involved. Collectively, our study identifies core transcriptional signatures associated with stem cell induction and elucidates the dynamic regulatory mechanism driving early stem cell lineage establishment.

## INTRODUCTION

Developmental plasticity in plants relies on their ability to reprogram and generate pluripotent and stem cells throughout almost their entire life cycle^1, 2^. This unique characteristic is intimately associated with plant postembryonic development. As the plant formative program is not completed during embryogenesis, subsequent organogenesis and growth is observed, necessitating the initiation, establishment and maintenance of new stem cell niches from non-pluripotent cells.

Stem cell niches are microenvironments where stem cells are located ^2, 3^. Stem cell niches support organ growth and tissue patterning and their activity and maintenance is tightly regulated by stem cell organizers. In roots, the stem cell organizer is known as the quiescent center (QC) because of the slow division rate, which provides signals for stem cell homeostasis, maintains a low differentiation rate, and replaces damaged stem cells during growth and regenerative processes^4, 5^. QC specification and homeostasis directly involves *WUSCHEL RELATED HOMEOBOX 5* (*WOX5*) ^6^, whose expression is regulated by the stem cell factors SHORT-ROOT (SHR), SCARCROW (SCR) and PLETHORA (PLT) 1 and 2, among others ^4, 6, 7, 8, 9, 10^. Remarkably, these stem factors are not only been associated with stem cell niche specification during growth^11^ but also with the formation of new stem cell niches during regeneration^12^. It has been proposed that full developmental plasticity allowing cell reprogramming (endogenously or upon hormone supplementation) to a pluripotent state, is only possible in cells carrying a pericycle-like identity^13, 14^. However, most meristematic cells surrounding stem cell niches can be reprogramed to form a new stem cell niche upon ablation or resection^15^, suggesting that the maintenance of certain meristematic feature underlies the process and that these features are kept in the pericycle^1, 16^.

Non-embryonically originated stem cell niches are also common in plants and associate with organogenesis processes during aerial and root system development. Tap root systems, as observed in *Arabidopsis thaliana*, develop through lateral root formation, a mechanism that naturally induces and establishes new stem cell lineages from specific subsets of pericycle cells^17, 18^. Remarkably, wounding and hormone-induced regeneration follow a lateral root developmental program^13, 16^, supporting the importance of investigating root postembryonic organogenesis and stem cell induction.

The recruitment of specific subsets of pericycle cells to form founder cells is key for lateral root organogenesis^19, 20, 21, 22^. Pericycle reprogramming competence, maintained by several bHLH transcription factors, is critical for founder cell specification^23^. It has been proposed that local accumulation of auxin^21^, activating the expression of certain factors such as GATA23, is also required in this process^24^. Root founder cells undergo a series of asymmetric divisions, a process morphologically preceded by nuclear migration^25, 26, 27^. As a result, 2-4 smaller cells in the central domain and two larger cells in the lateral domain are formed^22, 28^. Subsequently, a series of anticlinal, periclinal and tangential divisions generate a dome-like structure without a deterministic morphological structure but with a conserved tissue pattern^29, 30^. The process of lateral root formation has been categorized in seven developmental stages^17^. Lateral root tissue patterning involves factors such as PLT 3/5/7 activating PLT1 and 2, as well as stem cell regulators such as SCR, which is critical for initiating a new QC^31, 32^. The process culminates in later developmental stages with the activation of the new meristem and the outgrowth of the new organ^17^.

As new organs are formed, cell identity is primarily specified based on the origin of the cell (i.e., its lineage), its morphology, and its position within the organ, although additional features such as cell functionality or an altered molecular state also contribute^33^. Genome-wide expression data obtained through RNA sequencing (RNA-seq) or single cell RNA sequencing (scRNA-seq), often used to reconstruct gene regulatory networks (GRN), has been used to gain insight into formative processes, cell identity specification and stem cell homeostasis^34, 35, 36, 37^. RNA-seq of gravistimulated roots^38^ allowed for the reconstruction of a lateral root GRN^39^, showing the involvement of AUXIN RESPONSE FACTOR (ARF) 7 and ARF5 in the establishment of central and flanking developmental domains during lateral root formation through their mutual inhibition. This regulation could require additional modulation through Aux/IAA inhibitors to activate bi-modular auxin responses^40^, potentially contributing to organ symmetry. scRNAseq of gravistimulated roots captured specific gene upregulation associated with lateral root initiation and xylem pole pericycle cell reprograming^41^.

Specific scRNA-seq of the earliest lateral root formative stages has revealed a root ontological hierarchy in Arabidopsis^18^. Root organogenesis involves three developmental trajectories derived from primordial cells (i.e., the founder cells and their daughter cells) organized following 3 perpendicular developmental axes. The establishment of these developmental trajectories requires the specification of precursor cell types, which in the stem cell trajectory is particularly interlinked with stem cell induction. Genetic components play a major role in the specification of precursor cell types, as illustrated by analyses in the mutant of the C-REPEAT BINDING FACTOR 3^18^ or through the selective repression of target transcription factors in xylem pole pericycle cells^41^. Although all this evidence points to postembryonic root organogenesis relying on the generation of a complex network of positional and genetic regulations, the mechanism leading to the formation of stem cell precursor cell types and driving stem cell induction, remains unknown.

To further investigate postembryonic stem cell induction and precursor stem cell type specification, we reconstructed a spatiotemporal, cell-type-specific transcriptional map of early lateral root organogenesis in Arabidopsis. Using these data, we reconstructed a GRN in the early root stem cell lineage that reveals that stem cell initiation is driven by a morphogenic cascade of hierarchical interdependent transcription factors and identifies core transcriptional signatures associated with stem cell induction. Overall, our study elucidates the dynamic regulatory mechanism driving stem cell lineage establishment.

## RESULTS AND DISCUSSION

### Detailed Spatiotemporal Transcriptomics of the Early Stages of Lateral Root Organogenesis

Lateral root organogenesis is hierarchical^18^, imposing a developmental sequence that begins with the reprogramming of the specialized pericycle at the xylem pole ends^16, 41^. Thus, our experimental approach to capture the most relevant transcriptional changes associated with the initiation and onset of the new organ (Fig. S1a) started by profiling the pericycle, followed by the subsequent early generated populations: founder cells, QC/Endodermis transitioning cells and QC transitioning cells.

Specific fluorescent markers exist for pericycle tissues^42^<u>_ENREF_1</u>, so in order to explore the pericycle reprogramming capacity, we used a general pericycle marker (J0661, population 1) and a specific XPP marker (J0121, population 2) (Fig. 1a, b) to profile these cell types through Fluorescence Activated Cell Sorting (FACS; Fig. S2a, b). The XPP gives rise to founder cells, and thus, to capture this specification event, we analyzed in detail, through confocal microscopy, founder cell fluorescent markers previously generated by us^18, 43, 44^. We found that the SKP2B marker was associated with founder cell specification (population 3), being expressed earlier than the HB53 marker, which was associated with founder cell nuclear migration (population 4) (Fig. 1a, b). After careful examination of hundreds of plants under the confocal microscope, we determined that under our experimental settings, founder cells never divided earlier than 3 days post imbibition (dpi) (Fig. S1b), enabling specific FACS profiling of the founder cell types (Fig. S2c, d). In addition, we profiled a slightly later time (Fig. S1b) when founder cells had started to divide to capture gene expression associated with primordial cells (population 5, Fig. 1a, Fig S2c, d). Given asynchrony in the initiation of founder cell division, this sample contained a mixture of founder and stage I cells.

**Figure 1.**
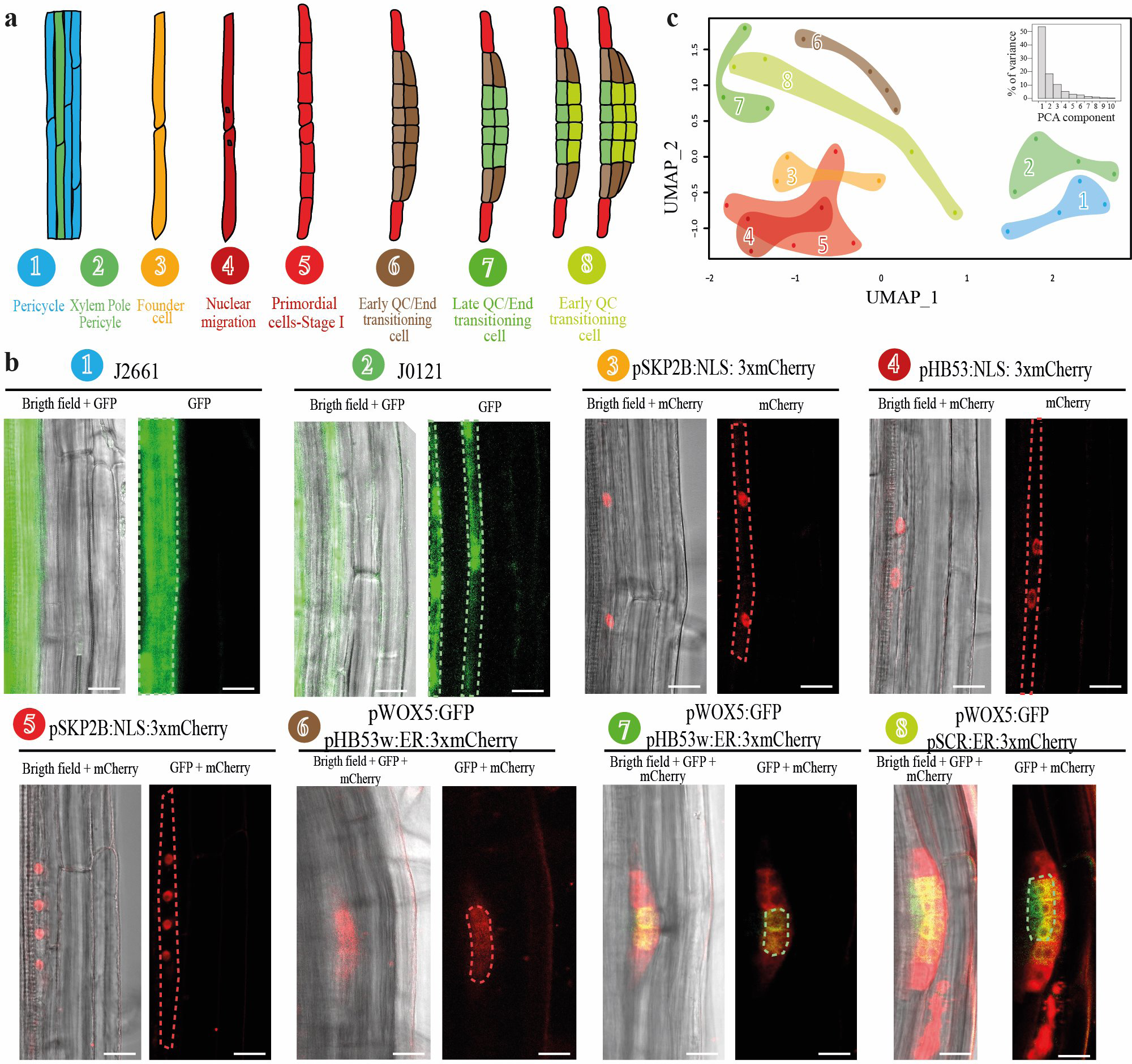
Profiling the Early Stages of Lateral Root Organogenesis. **a)** Schematic cartoon illustrating the cell types isolated during early lateral root formation through Fluorescent Activated Cell Sorting (FACS) and used for RNA sequencing (RNA-seq) transcriptomic analyses. **b)** Confocal laser microscopy images of the fluorescent marker lines used in this study. Left panel: overlapping bright field and fluorescence images. Right panel: fluorescence images. J0121 and J0661 are enhancer trap lines driving expression of the Green Fluorescent Protein (GFP), whereas the other lines correspond to transcriptional fusions of the promoters (p) of *HOMOLOG OF HUMAN SKP2.2* (SKP2B), *HOMEOBOX53* (HB53) and *SCARECROW* (SCR) to the mCherry fluorescent protein, or of *WHUSCHEL RELATED HOMEOBOX 5* (WOX5) to the GFP and their combinations as indicated. Scale bars: 5 µm. **c)** Principal Component Analysis and Uniform Manifold Approximation and Projection (UMAP) dimensional reduction of the transcriptomes 1 to 8 from a). Each dot corresponds to a biological replicate. UMAP utilizes PCA1 to 6 explaining >95% of variance. Color legend is as shown in a).

The HB53 marker is consistently expressed up to stage IV lateral root primordia, while the WOX5 marker activates during the formation of the new QC. In order to profile gene expression associated with QC/Endodermis transitioning cells, we generated double fluorescent markers by transforming the HB53-mCherry marker into plants carrying the WOX5-GFP marker. In one of the resulting lines, which we named WOX5/HB53w, the mCherry was only visible from stage II (Fig. 1a, b), allowing, in combination with the collection of samples at 4 dpi (Fig. S1b), when most lateral primordia were at stage II (∼90%) under our experimental settings, for the FACS profiling of samples highly enriched in QC/Endodermis transitioning cells. We collected this cell type before (population 6; Fig. S2c, d, please note the absence of GFP positive cells in the R8 area) and after the appearance of WOX5 expression (population 7; Fig. S2a, b). In our experimental setting we dissected the root maturation zone (Fig. S1a) to prevent contamination of samples with WOX5-marked QC cells from the root apical meristem, which is confirmed by not detecting GFP positive cells in population 6 (Fig S2d, R8 area, Ratio R8/R5<0.0018) although those plants expressed the WOX5 marker in the primary root apical meristem.

Finally, we transformed the SCR-mCherry marker into WOX5-GFP expressing plants to facilitate capturing the QC transitioning cells (Fig. 1a, b) through FACS, as those cells marked with both the mCherry and the GFP. Although the fluorescence intensity of these markers was unfortunately not bright enough to sort cells doubly marked with the two fluorescent proteins through FACS, we could successfully obtain a sample enriched in the early QC transitioning cells (population 8, Fig. 1b, Fig. S2a, b) by collecting samples through the WOX5-GFP marker at a later time, closer to 5 dpi (Fig. S1b), when there was a fair proportion (∼70%) of stage III lateral root primordia, yielding a greater proportion of GFP positive cells in population 8 (R2/R5 ∼ 3%) than in 7, (R2/R5 ∼ 0.5%).

Following FACS, we determined the transcriptomes of the 8 cell populations by RNA sequencing (RNA-seq; Table S1). Dimensional reduction of the transcriptomic data showed that related samples such as populations 1 and 2 (pericycle), populations 3, 4 and 5 (founder and primordial cells) and populations 6, 7 and 8 (QC/Endodermis- and Early-QC-transitioning cells) tended to cluster together (Fig. 1c). Two of the early QC transitioning cell samples showed more intermediate positions in the plot, which might correspond to intermediate developmental states towards fully specified QC cells and suggests the existence of distinctive transcriptomes for QC transitioning cells. Collectively, our time-course approach sampling cell-type specific fluorescent markers generated a unique dataset for early postembryonic organogenesis.

### The Lateral Root Organogenesis Transcriptional Map Establishes Novel Spatialtemporal Dynamics for Stem Cell Regulation

Previously described stem cell regulators were expressed through our postembryonic organogenesis dataset (Fig. S3). Notably, some of these factors, such as the PLTs and SCR, were also described to regulate lateral root formation ^31, 32^. Our analysis shows that *PLT3/5/7* peak at the earliest formed cell populations, while *PLT1/2/4* started to be expressed at late QC/Endodermis transitioning cells, which is in agreement with *PLT3/5/7* being expressed at stage I of lateral formation to regulate cell type pattern at later developmental stages through the activation of *PLT1/2/4*^31^. <u>_ENREF_9</u>Our data also show that *WOX5* starts to be expressed in the late QC/Endodermis transitioning cells, which very well matched our observation of the WOX5 marker appearance in this cell type (Fig. 1b). As PLT1/2 maintain *WOX5* expression and the stem cell niche of the primary root apical meristem^45^, it is possible that *WOX5* expression is maintained by PLT1/2 in QC/Endodermis transitioning cells, where all these factors concurrently peak (Fig. S3). Interestingly, SCR starts to be expressed in early QC/Endodermis transitioning cells preceding *WOX5* activation, which is also in agreement with SCR regulating QC initiation^32^. Interestingly, SCR, SHR, JKD, and TCPs, also peaked before *WOX5* activation, while they had been previously shown to regulate *WOX5* in the primary root^11^. Furthermore, *SHR* expression also preceded activation, in our dataset, of its well-known direct targets *SCR* and *JKD*^46^. Therefore, out early organogenesis transcriptional map not only recapitulates expression patterns which are in agreement with known regulatory interactions, but, most importantly, it establishes novel spatialtemporal dynamics associating stem cell regulators with precursor cell types at certain developmental times in the organogenesis stem cell lineage.

### Specific Transcriptomes are Associated with Organogenesis-Specific Cell Types and Their Developmental Progression

Our previous analyses suggest that a regulatory cascade could be established along a spatiotemporal developmental trajectory to initiates a new stem cell niche through the specification of precursor cell types such as the QC/Endodermis- and QC-transitioning cells. To investigate more in depth, the spatiotemporal developmental trajectory profiled by our early organogenesis dataset, we performed differential expression analysis of the samples following the order given by cell type formation (Fig. 2a). This approach would identify temporal gene enrichment or depletion during organogenesis. We discerned that the differentially expressed genes (DEG) between the pericycle (population 1) and the XPP (population 2) would relate to its reprogramming capacity, while the DEG between XPP (population 2) and founder cells (population 3) would capture founder cell specific characteristics. In addition, the DEG between founder cells (population 3) and founder cells during nuclear migration (population 4) would render enrichment in the asymmetric-division-related genes initiating organogenesis. As the primordial cell sample (population 5) also contained founder cells, we did not include it in the differentially expression analyses to more specifically identify genes primarily associated with temporal progression. The DEG between the subsequently formed early QC/Endodermis transitioning cells (population 6) and the nuclear-migration-state of founder cells (population 4) would provide the genes associated with this stem cell precursor cell type. Finally, the DEG between developmental stages intimately associated with *WOX5* induction or expression, such as the DEG between the late (population 7) and early (population 6) QC/Endodermis transitioning cells, and between the Early QC- (population 8) and late QC/Endodermis- (population 7) transitioning cells would inform about the acquisition of stem cell characteristics and genes involved in the developmental progression towards the formation of a new stem cell niche.

**Figure 2.**
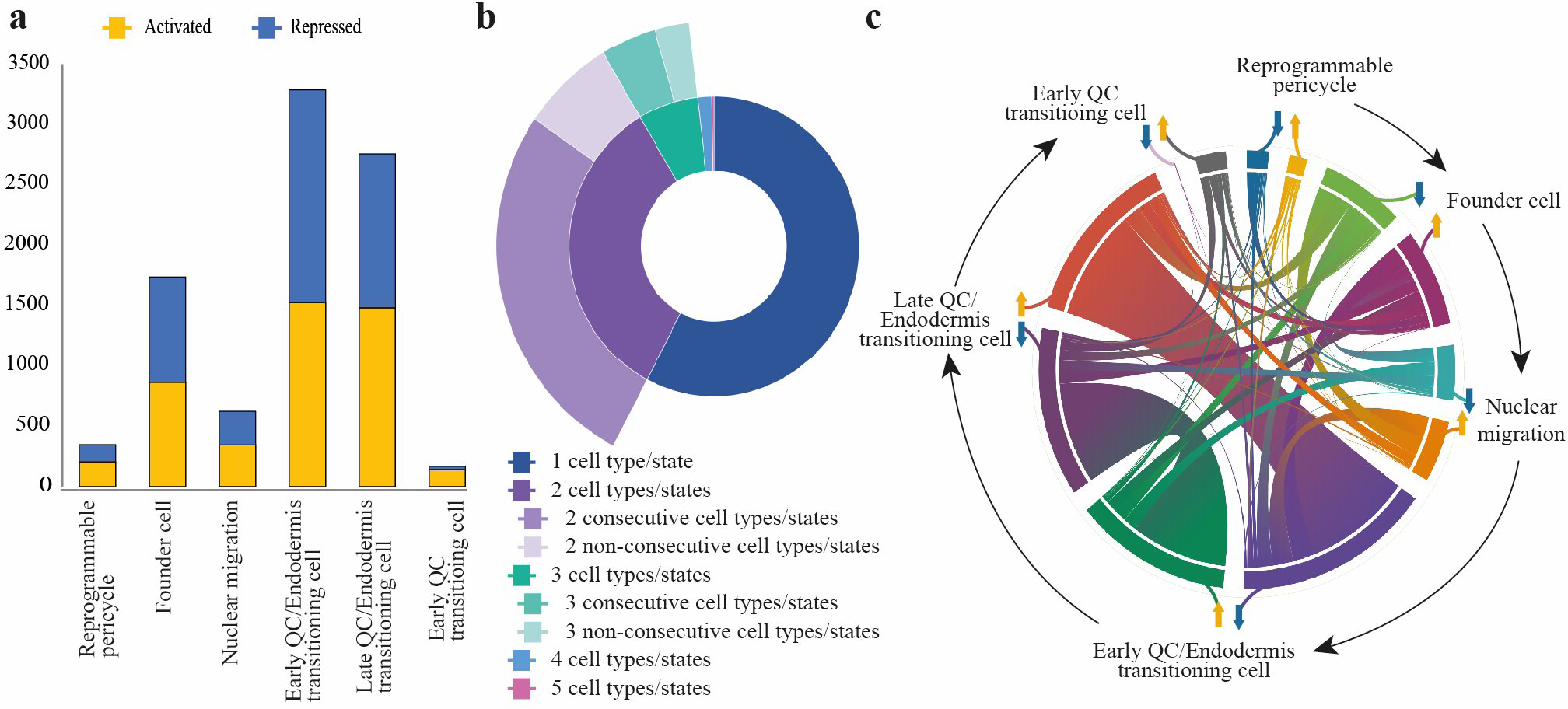
Profound Transcriptomic Changes Occur During Cell Reprograming and Early Organogenesis. **a)** Bar chart illustrating the characteristic number of activated (yellow) or repressed (blue) genes in various cell types and cellular states during the early stages of lateral root organogenesis. Reprogrammable pericycle: differentially expressed genes (DEGs) between samples 1 and 2 –as shown in Fig.1a-; founder cells: DEGs between samples 3 and 2; nuclear migration: DEGs between 4 and 3; early and later QC/Endodermis transitioning cells: DEGs between 6 and 4, and 7 and 6, respectively; and early QC transitioning cells: DEGs between 8 and 7. **b)** Sunburst chart depicting the proportion of DEGs present in one or more cell types or cellular states, and whether these correspond to temporally consecutive or non-consecutive cell types generated during organogenesis. **c)** Chord diagram illustrating genes induced (yellow arrows) or repressed (blue arrows) in various cell types and cellular states, with two main groups of genes switching their expression behavior (from activated to repressed and vice versa) between early and late QC/Endodermis transitioning cells at the onset of *WOX5* biomarker induction.

A total of 5832 DEGs were identified to dynamically change during all formative transitions (Fig. 2a, Table S2). DEGs in early and late QC/Endodermis transitioning cells were the most abundant, followed by DEGs in the founder cells, which could refer to their condition as newly formed cell types. In contrast, smaller numbers of DEGs were identified for reprogrammable pericycle, nuclear migration and early QC transitioning cells, which could be interpreted as certain cell states requiring less regulations to be initiated or maintained. 85% of the genes were differentially expressed in only one (3361 DEGs, 58%) or two consecutive developmental events (1570 DEGs, 27%), highlighting the specificity of our transcriptomic analyses (Fig. 2b). Only the remaining 15% of the genes were differentially expressed in more than two events, although yet one third of them were differentially expressed sequentially. No DEGs were detected across all the six comparisons. Our analyses have identified DEGs as proxy for gene enrichment and depletion across the 6 cell types or states, and as a result of their specificity and dynamics these DEG could not only relate with specific cell-type characteristics but also participate in the underlying formative events.

### Profound Transcriptomic Changes Occur During Formation of Organogenesis Specific Cell Types

To further investigated which cell types or states shared DEG and their distribution we performed a multiple Venn Diagram analysis. Remarkable, most of the DEG were shared across sequentially formed cell types or states (Fig. S4), thus they could function as connectors, reinforcing the idea of a regulatory cascade propagating stem cell induction along the stem cell trajectory. In agreement with only a few proportion of the DEG being shared by more than two cell types or states, as previously shown (Fig. 2b), we only observed a few tens of genes not being shared sequentially (Fig. S4).

We performed a chord diagram for the DEG to investigate the connections between the genes induced or repressed across the different formative events (Fig. 2c). We observed two main flows of genes corresponding to those induced and them repressed in the early and late QC/Endodermis transitioning cells, respectively, and between those initially repressed and then activated in the same cell types. This reversion from activated to repressed and vice versa suggests a profound transcriptomic rearrangement during the specification of the precursor lineage giving rise to a new stem cell niche. A comprehensive transcriptional signature of stem cell induction is lacking^35, 36, 47, 48^, our specific transcriptomic data can shed light into the transcriptomic drivers of stem cell induction, therefore, contributing to define stemness.

### Comprehensive Inference of Mechanistic Regulation during Early Organogenesis and Cell Type Specification

Gene regulatory networks (GRN) represent regulatory interactions among sets of genes and can predict dynamic spatialtemporal regulation ^35, 36, 49^. We used our transcriptomic dataset to infer spatial and temporal interactions among the genes involved in stem cell induction during organogenesis. By applying a dynamic Bayesian network (DBN) inference algorithm, genist, and a regression tree-based pipeline, rtp-star, we determined transcription factor regulatory dynamics and downstream regulation (Fig. 3a). This network recapitulated the specification of 5 precursor stem-cell types or states in the stem cell lineage, including founder cell specification as a landmark for pluripotency (Fig. 3b-f). As transcription factors (TFs) integrate positional and temporal information we focused on the 377 DEG TFs as drivers of stem cell induction (Fig. 4a). Analyses of enrichment in Gene Ontology (GO) categories for these DEG TFs showed significance in biological processes such as cell differentiation, regulation of developmental process, pattern specification process, tissue development or response to auxins and other hormones (Table S3).

**Figure 3.**
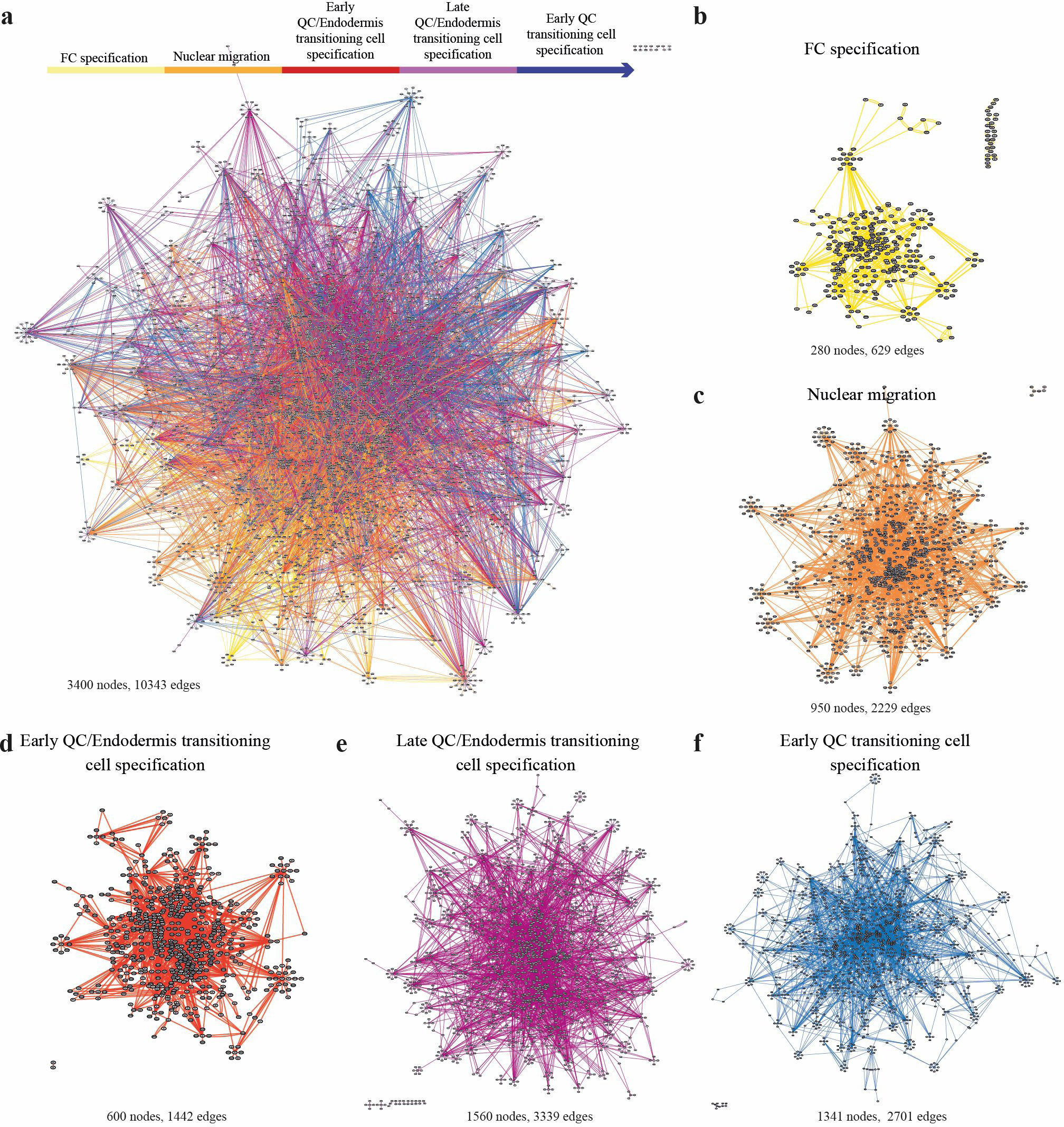
A Gene Regulatory Network for Early Organogenesis and Cell Type Specification Predicts Mechanistic Regulation. **a)** A temporal gene regulatory network (GRN) was reconstructed through the GENIST Bayesian network inference algorithm from the DEG TF to infer TF-TF regulation, and combined with the rtp-star regression tree pipeline from all the DEG to infer TF-gene regulation in the cell types and cellular states profiled during the early stages of lateral root organogenesis. Five different developmental transitions were defined as depicted in the legend. **b-f)** Subnetworks for the five early organogenesis transitions predict cell fate or state specification events.

**Figure 4.**
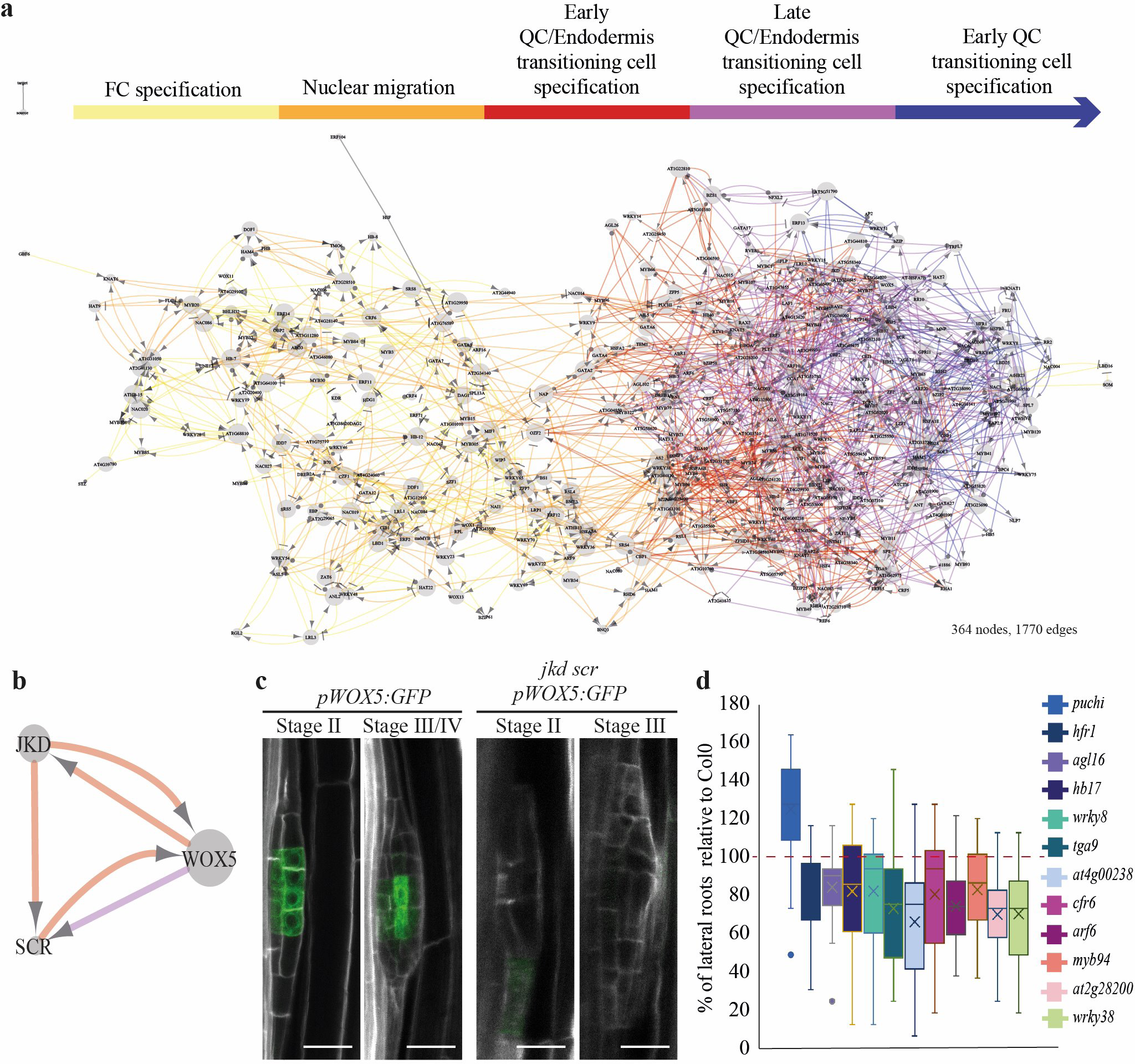
Organogenesis Progression Is Driven by a Hierarchical Network of Sequentially Interconnected Transcription Factors. **a)** A temporal Gene Regulatory Network (GRN) for the DEG TF (out of the TF-TF-gene GRN of Fig. 3) in cell types and cellular states profiled during the early stages of lateral root organogenesis is shown. Five different developmental transitions were defined, as depicted in the legend. **b)** Subnetwork showing regulation among *WOX5*, *JKD* and *SCR*. Arrow colors correspond to those in a). **c)** Confocal laser microscopy images showing expression of *pWOX5:ER-GFP* in the wild type or in *jkd scr* during the first stages of lateral root organogenesis. Scale bars: 25 µm. **d)** Fifty TF were randomly selected from those most highly connected in the GRN, and mutants in those genes tested for lateral root capacity. The boxplot shows the ratios mutant/control with statistically significant changes: p-value < 0.05 in a Univariate General Linear Model; n>25 in two biological replicates.

### Stem Cell Initiation is Driven by a Morphogenic Cascade of Hierarchical Interdependent Transcription Factors

Most of the TFs (83%) in the early organogenesis GRN connected two consecutive specification events. A more detailed analysis of the dependencies in the GRN (Fig. S5a), showed sequential regulatory interdependency, where the specification of a cell type or state triggered initiation of the subsequent one in the lineage. Thus, a fair number of TFs involved in founder cell specification also regulate nuclear migration and thus, a sequential regulatory cascade is triggered determining progression across the 5 formative processes once the first one is started. In this GRN structure, each formative stage could require completion of the previous one for developmental progression, which could be delivered by the TF specifically associated with each one of the five specification events. Therefore, this GRN structures as a morphogenic cascade, which in animals normally associate processes triggering cell fate determination and difficult to reverse^50, 51, 52^. Considering the flexibility observed in plant development, it would be interesting to explore if fate could be reversed along the stem cell initiation trajectory. However, our previous laser ablation experiments on root precursor cell types, which appeared to restart the sequence of identity initiation in the stem cell trajectory, suggest irreversibility^18^. It was notorious the high number of TFs (64%) participating in QC/Endodermis transitioning cell specification, in which most of the regulators (77%) where shared among the early and later developmental stages. Finally, we compared our GRN with other networks involved in wound/hormonal induced reprogramming and stem cell ubiquitous regulation showing high specificity although some transcriptional signatures were observed (Fig. S5b). It is possible that these commons transcriptional signatures associate with pluripotency or stemness acquisition.

Certain network motifs, such a feedback and feed-forward loops, have been previously shown to be enriched in GRN associated to cell fate and stemness regulation^35, 36, 51, 53^. Among the multiple cases of these types of regulatory connections in our GRN, the feedback regulation among WOX5, JKD and SCR during QC/Endodermis transitioning cell specification (Fig. 4b) particularly drew our attention. SCR and JKD had been previously shown to regulate *WOX5* expression and stem cell niche specification in the primary root^11^; and, in addition SCR regulates QC initiation and regeneration ^12, 32, 46^. Confocal microscopy analyses of the double mutant *scr jkd* showed impaired activation of *WOX5* during QC/Endodermis transitioning cell specification, along with obvious morphological defects (Fig. 4c). This observation together with the fact of stem cell transcriptomic signatures, such as *WOX5* and *PLT1/2* (Fig. S3), being initiated in QC/Endodermis transitioning cells indicates that this cell type can be considered a stem cell progenitor and be key for stem cell induction.

We leveraged the topology of our GRN to identify key regulators based on a higher degree of connectivity (edgecount>10). We selected a total of 50 TFs and perform a lateral root capacity assay upon homozygous mutant lines to assess their involvement in lateral root organogenesis. We found 11 TFs mutants with a significantly reduced number of lateral roots, while one, PUCHI, displayed increased number (Fig. 4d). Interestingly, the mutant *puchi* was previously shown to have increased lateral root number (Kang et al., 2013). These results confirm the power of our approach and GRN to identify novel mechanistic regulation during early root organogenesis and cell type specification.

## EXPERIMENTAL PROCEDURES

### Plant material and growth conditions

*Arabidopsis thaliana* Columbia-0 (Col-0) accession was the genetic background used in this study. Seeds were surfaced-sterilized upon exposition to chloro gas and stratified in sterile water at 4°C in darkness during 2 days. After stratification, seeds were transferred to petri dishes containing half-Murashige & Skoog (MS) medium with 1% sucrose and 10 g/L Plant Agar (Duchefa). Arabidopsis seedlings were vertically grown in chambers under a 16/8 photoperiod at 22°C. J0121^42^, J2661^42^, pWOX5::ER-GFP^6^, pHB53:NLS-3xmCherry^18^ and pSKP2Bs::NLS-3x-mCherry^43^ lines were used in this study. *dpGreen BarT pSCR:ER:3xmCherry* construct was generated using MultiSite Gateway Three-Fragment Vector Construction Kit (Invitrogen) from previously generated pDONOR plasmids^18, 54^ and together with the *dpGreen BarT pHB53::NLS-3x-mCherry* contruct^18^ introgressed into the pWOX5::ER-GFP line^6^ using the floral dip method to generate the pHB53w::NLS-3x-mCherry pWOX5::ER-GFP and pSCR::NLS-3x-mCherry pWOX5::ER-GFP lines. Homozygous lines were selected among the T2 progeny in ammonium glufosinate (Merck). The mutant *jkd scr* pWOX5::ER-GFP line was generated by multiple crossing of *jkd-4* and *scr-4* mutants with pWOX5::ER-GFP followed by genotyping through PCR using previously described primers^46^ and analysis of fluorescence segregation (see Microscopy and Imaging analyses) to identify stable homozygous lines in the F3 generation. Rest of mutants in this study are those listed in Table S4. Homozygosity was confirmed using the generic primers obtained through T-DNA Primer Design (http://signal.salk.edu/tdnaprimers.2.html).

### Protoplast isolation and RNAseq

The biological samples were extracted from plants expressing the different marker lines as indicated. For each independent replicate, the primary root tip and the aerial part of ∼400 plants were removed as indicated (Fig. S1) under a fluorescent dissecting scope and the remaining roots were subjected to 1.5 hours of protoplasting^55^. Each of the cell types with positive GFP/mCherry expression (Fig. S2) were collected by FACS (Beckman Coulter MoFlo XDP) in 15 mL tubes (Falcon) with 3 mL of QIAGEN RLT buffer and 30 μL of β-mercaptoethanol, to minimize messenger RNA degradation. Between 5-6 bilogical samples were collected per sample. The commercial RNeasy Micro Kit (QIAGEN) was used to extract messenger RNA from the cells of interest isolated by FACS, according to the manufacturer’s specifications and SMART-Seq v4 Ultra Low Input RNA Kit (TAKARA) was used to synthesize cDNA from messenger RNA. cDNA fragmentation was performed in COVARIS S220 according to previously established technical parameters^35^ to obtain an average size of 500 base pairs. Libraries were synthesized using the TAKARA commercial library preparation kit (Low Input Library Prep Kit v2), according to the kit specifications. Sequencing method used was single end sequencing of 100 base pairs on the Illumina HiSeq 2000 sequencer. The data was deposited in GEO under the accession number GSE244658.

### Analysis of massive sequencing data and dimensional reduction

We used the Tuxedo method^56^ within the Tuxnet pipeline^49^ for gene expression analysis. Briefly, samples were aligned to the Arabidopsis genome under standard conditions, the Cufflinks program was used to assemble the transcripts for each replicate according to the reference genome, and then the replicates for each sample were combined with the Cuffmerge program that allows for the final assembly of the transcriptome and calculation of differential expression fusing the Cuffdiff program. The expression value for each gene was normalized in FPKM (Fragments Per Kilobase Million). Additionally, we used the counts for each replicate prior differential expression analyses to perform dimensional reduction by Principal Component Analysis (PCA) and Uniform Manifold Approximation and Projection (UMAP) dimensional reduction using “princomp” and “umap” functions in in R, “version 4.3.2 (2023-10-31 ucrt)”, (https://www.R-project.org/). UMAP was performed using first 6 PCAs that accounted for >95% of variance.

### Bioinformatics Multiple Comparison analyses

Multiple comparison analyses of differentially regulated genes and flow of regulation were performed using the available on line tools “Calculate and draw custom Venn diagrams” (http://bioinformatics.psb.ugent.be/webtools/Venn/) and Venny 2.1 (https://bioinfogp.cnb.csic.es/tools/venny/). Chord diagrams were performed using the “chordDiagram” function in R, “version 4.3.2 (2023-10-31 ucrt)”, (https://www.R-project.org/).

### GO Enrichment analysis

GO enrichment analyses were performed in g:Profiler ^57^ using a p-value < 0.001 to detect the GO terms enriched for the DEG across early organogenesis.

### Reconstruction of Gene Regulatory Networks (GRN)

To infer the GRNs, we used the GENIST and rtpstar programs^35, 36, 49^<u>_ENREF_15</u> in MATLAB software. First, we inferred a Bayesian Dynamic GRN through GENIST using the transcription factors with differential expression between transcriptomes and used them in combination with their expression across the different samples organized following the collection time to establish a temporal order in development. Second, we used the rtpstar tree-based method transition by transition through 100 iterations to identify TF downstream regulation. These regulatory interactions were combined with the Bayesian Dynamic GRN to reconstruct a unified GRN. Cytoscape software^58^ was used to visualize GRN and analyze GRN parameters.

### Lateral root capacity assay

TF hubs were selected from those induced at any transition in the GRN and with a high degree of connectivity (edge count higher than 10 with any other TF). Homozygous T-DNA insertional mutant lines for these TFs were ordered (Table S4) and confirmed for homozygosity (see “Plant material and growth conditions”). Lateral root capacity assay was performed as previously described^59^.

### Microscopy and Imaging analyses

2-5 days post imbibition seedlings, as indicated, from homozygous lines were stained with 10 mg/mL Propidium Iodide (PI) (Sigma-Aldrich) or 1 mg/mL SCRI Renaissance 2200 dye (SR2200) (Renaissance Chemicals) and imaged in a Leica SP8 laser-scanning confocal microscope (Leica) using a hybrid detector counting mode and the following settings: GFP (excitation 488 nm, acquisition 505 - 535 nm), mCherry (excitation 561 nm, acquisition 600 - 650 nm), SR2200 (excitation 405 nm, acquisition 430 - 450 nm) and PI (excitation 561 nm, acquisition 600 - 630 nm). Fluorescence microscopy was performed in Leica M205FA adapted with Hamamatsu EMCCD X2 camera.

### Statistical analysis

Statistical differences were detected using R. Homoscedastic groups were analyzed using Analysis of variance (ANOVA) and Tukey HSD or Bonferroni post-hoc tests for one or more factors, respectively. Significant differences were collected with 5% level of significance.

## AUTHOR CONTRIBUTIONS

Conceptualization, M.A.M.-R., R.S., and J.C.; Methodology, M.A.M.-R., R.S., J.C., A.S.C. and P.P.-G; Formal Analysis, A.S.C., A.d.L.B., J.C., L.S.-R., C.d.B and M.A.M.-R.; Investigation: J.C., A.S.C., A.d.L.B., C.d.B and M.A.M.-R.; Resources, M.A.M.-R., R.S., P.C. and P.P.-G.; Writing-Original Draft, M.A.M.-R., J.C. and C.d.B; Writing-Review&Editing, M.A.M.-R., J.C., C.d.B. and R.S.; Supervision, M.A.M.-R., R.S.; Funding acquisition, M.A.M.-R., R.S. and P.P.-G.

## ACKNOWLEDGMENTS

This work was funded by Ministerio de Ciencia e Innovacion MCIN/AEI/10.13039/501100011033 and by “ERDF A way of making Europe” through grants PID2019-111523GB-I00 and PID2022-140719NB-I00 to M.A.M.-R., and by the ‘Severo Ochoa Program for Centres of Excellence in R&D’, grants SEV-2016-0672 2017-2021 and CEX2020-000999-S to M.A.M.-R and P.P.-G. through CBGP. A.S.-C. was supported by a FPI contract from MICIN (BES-2014-068852), P.P.-G. by a Juan de la Cierva contract from MICIN (FJCI-2015-24905) and Programa Atraccion Talento from Comunidad Madrid (2017-T2/BIO-3453) and J.C by a Juan de la Cierva contract from AEI (FJCI-2016-28607) from MICIN. The authors declare that they have no other competing interests.

---

**Figure S1.**
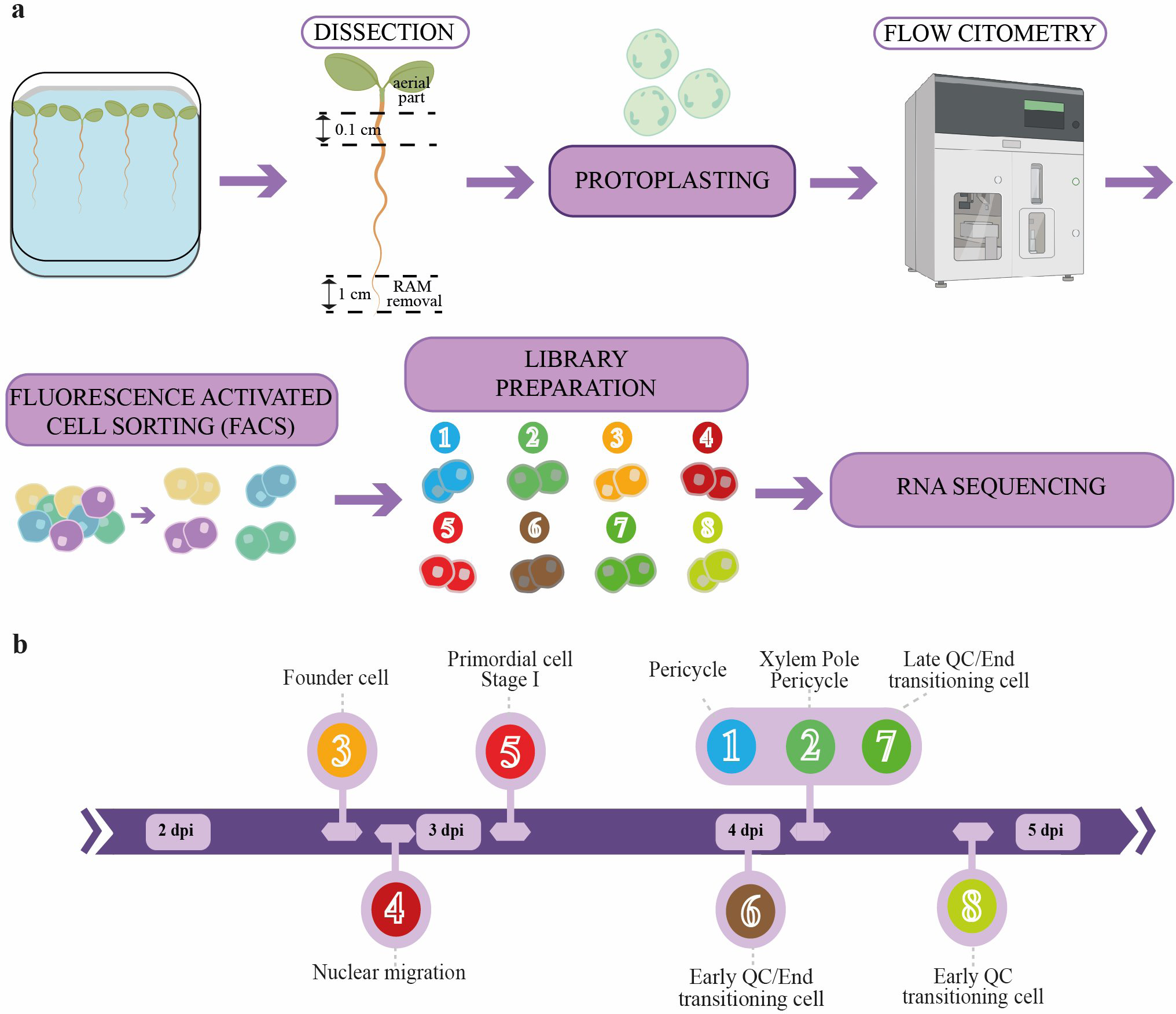
Flowchart Illustrating the Experimental Design for Profiling the Early stages of Lateral Root Organogenesis. **a)** The maturation zone of roots of the markers lines from Fig. 1c was dissected under a stereomicroscope to ensure no root apical meristems (RAMs) or aerial parts were collected. Samples were protoplasted and processed through Fluorescence Activated Cell Sorting (FACS) to isolate the various cell populations previously described. These sorted cells were used to prepare libraries and perform RNA-seq. b) Timeline for sample collection, as previously identified through confocal laser microcopy, containing the desired cell populations. At these times, no subsequent developmental stages of lateral root development were found. This, in combination with the indicated marker lines, allowed for specific cell type isolation. dpi: days post imbibition. Numbers and colors correspond to those indicated in Fig. 1a.

**Figure S2.**
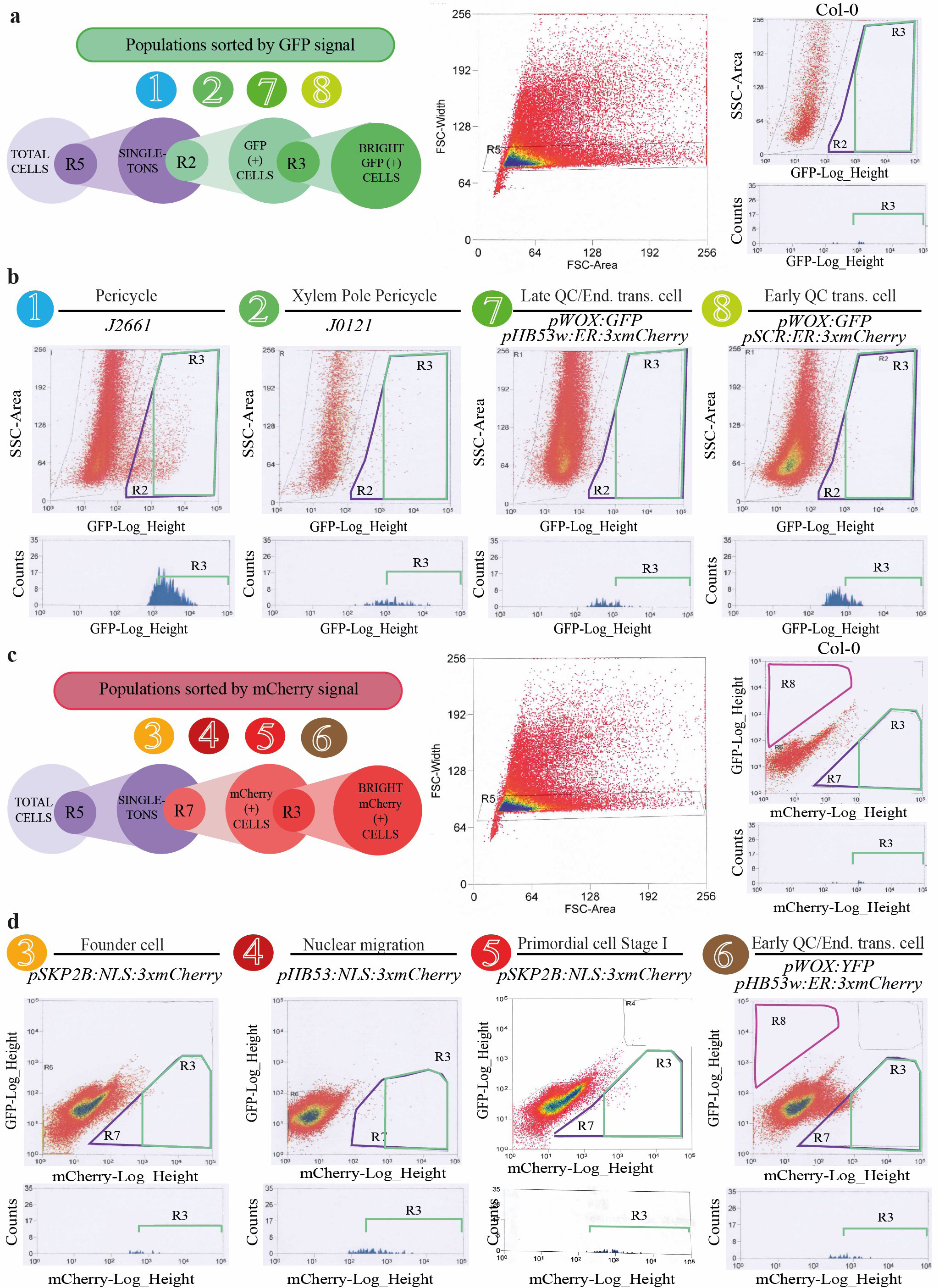
FACS of the Marker Lines Used for Profiling the Early Stages of Lateral Root Organogenesis. **a)** Schematics of the FACS setup used for sorting GFP-positive cells from marker lines and the outcome when applied to a control line without GFP-marked cells. **b)** Results obtained from applying the FACS setup previously described to the marker lines used to sort cell populations marked with the GFP. Only highly confident GFP-positive cells from the R3 region were used for transcriptomic analyses. **c)** Schematics of the FACS setup used for sorting mCherry-positive cells from marker lines and the outcome when applied to a control line without mCherry-marked cells. **d)** Results obtained from applying the FACS setup previously described to the marker lines used to sort cell populations marked with the mCherry. Only highly confident mCherry-positive cells from the R3 region were used for transcriptomic analyses.

**Figure S3.**
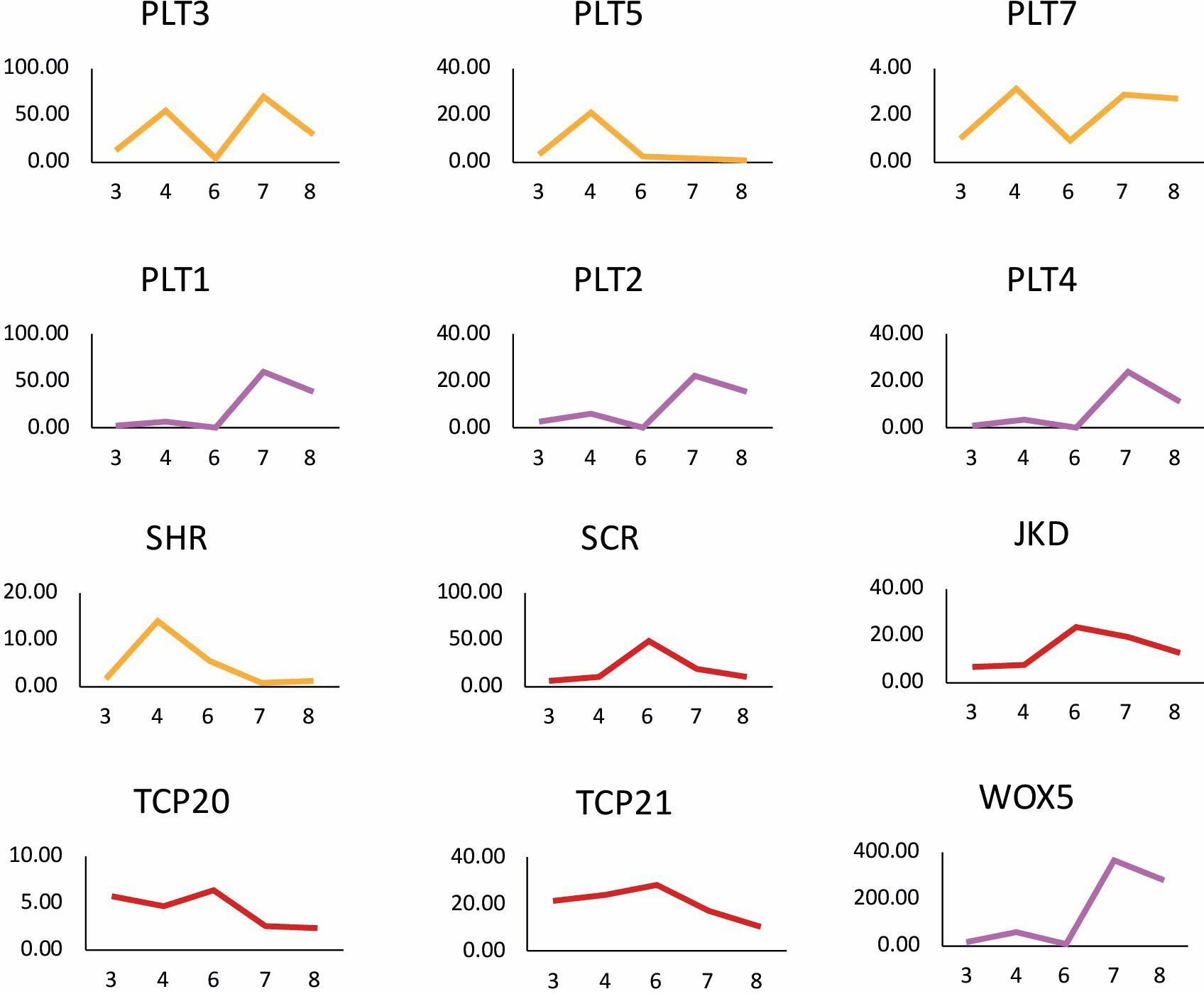
Expression Dynamics of Known Stem Cell and Lateral Root Formation Regulators. Graphs showing the expression of early-expressed regulators (PLT3/5/7 and SHR) in orange, mid-expressed regulators (SCR, JKD, TCP20 and 21) in red, and late-expressed regulators (PLT1/2/4 and WOX5) in purple across the founder cell (samples 3 and 4), QC/Endodermis transitioning cell (samples 6 and 7) and QC transitioning cell (sample 8) populations. Expression values are shown in Fragments Per Kilobase Million (FPKM).

**Figure S4.**
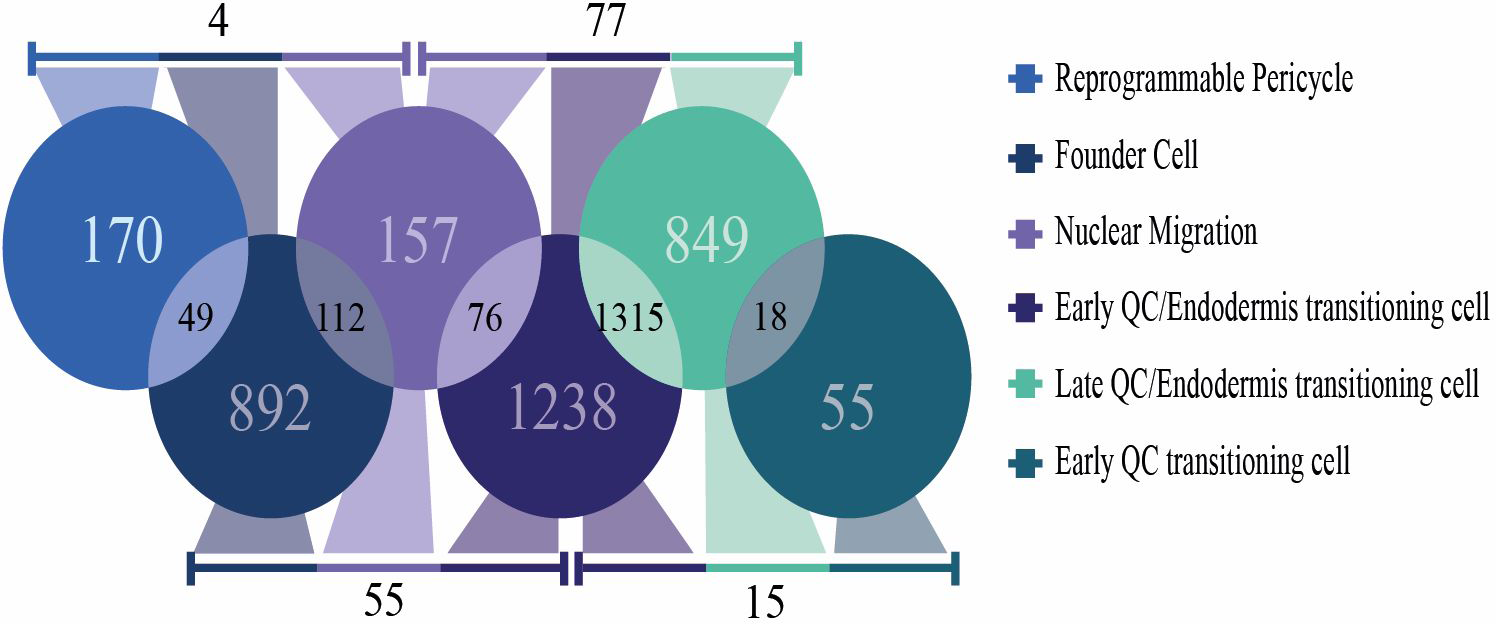
Specific Transcriptomes are Associated with Organogenic Cell Types and Their Developmental Progression. Venn diagrams show the number of DEGs present in a specific cell type or cellular state, or in common between two (numbers between circles) or more (numbers on the lines) of these cell populations.

**Figure S5.**
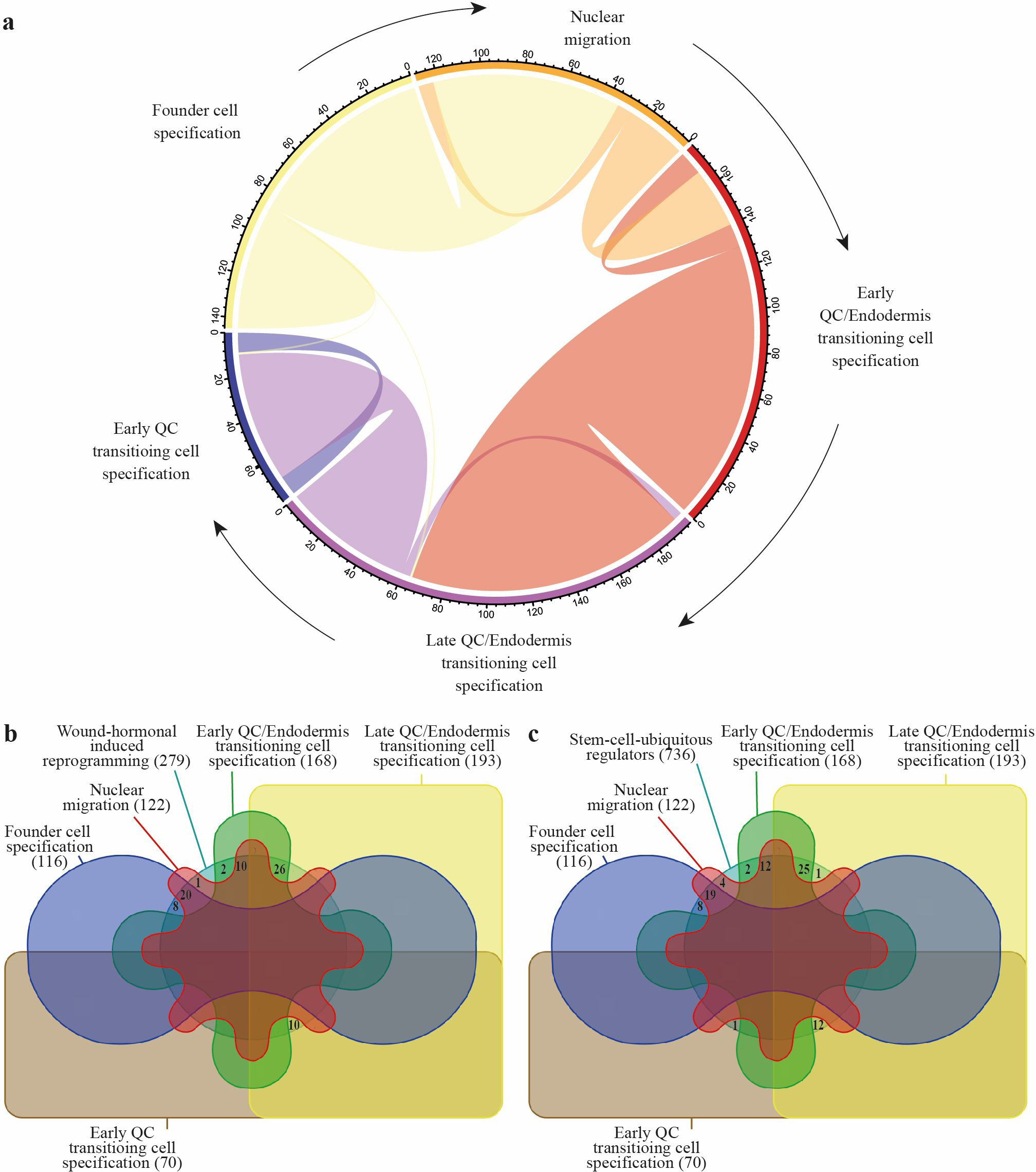
Stem Cell Induction is a Sequential Interdependent Process and Shares Certain Transcriptional Features with Reprogramming and Regeneration Processes. **a)** Chord diagram showing shared GRN TFs among the different developmental transitions. Colors correspond to regulatory interactions in GRN of Fig. 4a. **b, c)** Venn diagram showing common TFs among the GRN for the early organogenesis developmental transitions and GRN defined for b) wounding and hormonal reprogramming or c) stem ubiquitous regulation.

